# Immune protection and disease tolerance in *Pseudomonas aeruginosa* infections by pyruvate cross-feeding

**DOI:** 10.64898/2025.12.04.692321

**Authors:** Stefanos Lampadariou, Pablo Laborda, Lorenzo Masiero, Søren Molin, Helle Krogh Johansen, Ruggero La Rosa

**Affiliations:** The Novo Nordisk Foundation Center for Biosustainability, Technical University of Denmark, 2800 Kgs. Lyngby, Denmark; Department of Clinical Microbiology 9301, Rigshospitalet, 2100 Copenhagen Ø, Denmark; Department of Health Technology, Technical University of Denmark, 2800 Kgs. Lyngby, Denmark; Department of Clinical Medicine, Faculty of Health and Medical Sciences, University of Copenhagen, 2200 Copenhagen, Denmark

## Abstract

Chronic infections by *Pseudomonas aeruginosa* in people with cystic fibrosis are characterized by persistent inflammation and oxidative stress, yet the mechanisms enabling bacterial persistence are not fully understood. Here, we identify a novel persistence mechanism mediated by pyruvate secretion resulting from mutations in the pyruvate dehydrogenase complex in clinical isolates of *P. aeruginosa*, with analogous mutations also identified in *Staphylococcus aureus*, *Haemophilus influenzae* and *Stenotrophomonas maltophilia*. These mutations lead to elevated extracellular pyruvate which helps the establishment a more tolerogenic infection microenvironment and favor bacterial persistence. Pyruvate exerts multiple roles: scavenges reactive oxygen species such as H₂O₂, suppresses host immune activation both in airway epithelial cells and macrophages, and increases bacterial survival during phagocytosis. This metabolic crosstalk promotes bacterial persistence while limiting host tissue damage, aligning with the concept of disease tolerance. Our findings reveal pyruvate as a bacterial immunometabolite that mimics host antioxidant defenses, reshaping the infection niche to favor long-term colonization. This work highlights the broader role of secreted metabolites in host-pathogen interactions and suggests new strategies targeting metabolic pathways to manage chronic infections.

## Introduction

Metabolism, the intricate network of biochemical reactions within a cell, is the foundation of life. Far from being a passive background process, metabolism actively supports every single cellular function, ensuring the constant supply of energy and building blocks necessary for growth ^1^. In bacterial pathogens, metabolism plays an additional crucial role, directly influencing the ability to cause disease ^2,3^. Indeed, metabolic pathways are increasingly recognized as key players in the establishment and progression of bacterial infections by controlling host damage and its immune response ^4,5^. Strikingly, the precise and direct connections between specific metabolic pathways, the multifaceted mechanisms of bacterial pathogenicity and the host response often remain elusive ^6^.

*Pseudomonas aeruginosa* is an opportunistic pathogen causing both acute and chronic infections in several immunocompromised patients ^7^. Its ability to catabolize many carbon sources and colonize diverse environments and hosts reflects its extensive metabolic flexibility ^8^. Interestingly, it has been shown that adaptive evolution within the airway of people with cystic fibrosis (pwCF) selects for specific metabolic features which provide increased fitness during the infection ^9^. However, while the metabolic changes have been extensively described, the consequences of such metabolic specializations are largely unknown. One emerging hypothesis is that metabolic specialization selects for host-pathogen crosstalk leading to disease tolerance ^10^. The ability of the host to limit the damage caused by pathogens without decreasing the bacterial load, defined as disease tolerance ^10^, could actually be triggered by specific mechanisms of immune modulation employed by bacterial pathogens. For example, the immunometabolite itaconate mediates disease tolerance in *P. aeruginosa* pneumonia infection models by simultaneously inducing biofilm formation through alginate biosynthesis and limiting excessive inflammation by pulmonary glutamine assimilation ^4,11^. This and other mechanisms of host-pathogen metabolic interactions could, therefore, constitute broader mechanisms of protection, benefiting both the host and the pathogen ^12^. This is even more relevant in persistent infections where the outcome is the maintenance of the infecting population for long term rather than killing the host ^13^.

In pwCF, chronic *P. aeruginosa* infection is associated with sustained oxidative stress and excessive inflammation, primarily driven by reactive oxygen species (ROS) produced by neutrophils and macrophages. This process contributes to substantial tissue damage and progression of lung disease. Antioxidants are, therefore, necessary to maintain homeostasis and reduce inflammation. It has been proposed that the metabolite pyruvate, or its more stable form ethyl-pyruvate, elicits an antioxidant effect in several diseases due to its ability to react with H_2_O_2_ ^14–16^. Moreover, it has been described that ethyl-pyruvate down-regulates the production of pro-inflammatory cytokines such as interleukin-8 (IL-8) ^17^. Interestingly, in several bacterial species pyruvate has been described as a metabolic intermediate involved in pathogens protection from H_2_O_2_ and virulence regulation ^18–23^. However, whether pyruvate produced by bacterial pathogens modulates the immune system response and reduces the oxidative stress to promote persistence is unclear. We hypothesize that this persistence mechanism could influence disease tolerance, i.e. the ability of cells and tissues to limit host damage in the presence of invading pathogens, through a mechanism of metabolic crosstalk and host immunomodulation. This would help the establishment of a slow growing bacterial population, such as that present in pwCF, maintain tissue integrity and control organ damage.

Here, we provide evidence that pyruvate dehydrogenase complex (PDHc) mutations lead to high secretion of pyruvate which in the host microenvironment elicits a triple role. First, pyruvate directly reacts non-enzymatically with ROS species such as H_2_O_2_, protecting the bacterial community from oxidative killing. Secondly, it influences the host response by limiting IL-8 secretion by the airway epithelial cells, suppressing immune system cells recruitment. Third, it limits macrophages activation and increases bacterial survival within the phagolysosome. Indirectly, these mechanisms favor a less oxidative and more tolerogenic environment leading to disease tolerance by limiting airway inflammation. This is supported by the broad distribution of PDHc mutations in several clinical isolates of *P. aeruginosa*, *Staphylococcus aureus*, *Haemophilus influenzae* and *Stenotrophomonas maltophilia*. Altogether, these results demonstrate that persisting bacteria within the host select for PDHc mutations to influence disease progression through a generalized mechanism of metabolic crosstalk.

## Results

### Pyruvate dehydrogenase mutations protect against H_2_O_2_ stress

We hypothesized that mutations in the *P. aeruginosa aceEF* operon, encoding the PDHc, arising during chronic infections in pwCF ^24^ lead to pyruvate secretion as a protection mechanism against ROS due to the ability of pyruvate to react with H_2_O_2_ ^25–27^. To assess this hypothesis, we employed, along with the laboratory reference strain PAO1, two mutant strains, one harboring a point mutation in the *aceE* gene, leading to a Phe → Ser amino acid change at position 184, and the other one a 4 bp insertion in the *aceF* gene leading to a frameshift starting from Lys 273 ^24^. As previously reported, the *aceE* and *aceF* strains secrete pyruvate during growth in the defined medium SCFM2 which recreates the metabolic composition of sputum samples from pwCF and which does not contain pyruvate. Additionally, these strains show a fitness cost of the mutations underlined by their reduced growth rate ^24^ .

To assess the neutralizing reactivity of pyruvate against H_2_O_2_, we performed a H_2_O_2_ killing assay using an Air-Liquid Interface (ALI) model system consisting of mucociliated, differentiated airway epithelial cells that mimic the human airway environment. This system allows for the investigation of host responses, including epithelial integrity and immune system recruitment during infection. ALI models were infected with PAO1 or the *aceF* mutant strain for 6 hours to allow for the establishment of a bacterial population, followed by the addition of H_2_O_2_ on the apical side and incubated for an extra 30 minutes in presence or absence of pyruvate. Bacterial colony forming units (CFUs) from the planktonic fraction (apical compartment) and those attached to the epithelium were quantified to verify the bactericidal activity of H_2_O_2_ and the protective effect of pyruvate (Fig. 1A). As hypothesized, the addition of H_2_O_2_ significantly reduced the CFUs in both fractions, while the addition of pyruvate provided complete protection of the bacterial population (two-way ANOVA with Tukey’s multiple comparisons test, p<0.05).

**Figure 1.**
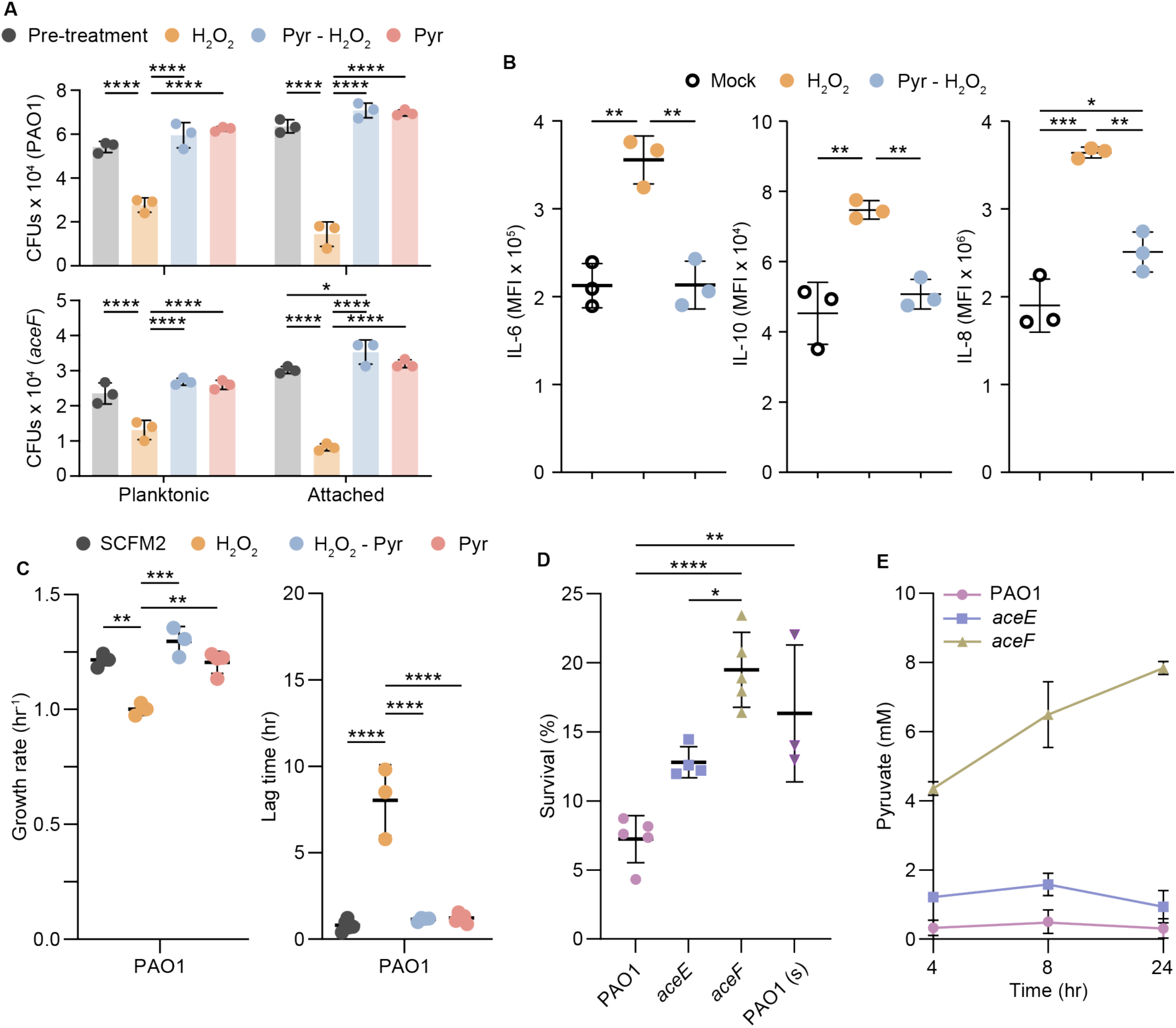
Neutralizing effect of pyruvate on H₂O₂ in *in vivo*-like models and laboratory conditions. (A) Colony-forming units (CFUs) of PAO1 and *aceF* mutant strain before and after H₂O₂ exposure in air-liquid interface (ALI) infection models of fully differentiated BCi-NS1.1 cells, with external supplementation of 2 mM H₂O₂ and 10 mM pyruvate. CFUs were quantified both for the planktonic bacteria in the apical compartment and attached to the epithelium. Statistical significance was determined using two-way ANOVA with Tukey’s multiple comparisons test. Significance is indicated as * (p < 0.05), ** (p < 0.01), *** (p < 0.001), and **** (p < 0.0001). (B) Cytokines release upon oxidative stress induction with 5 mM H₂O₂ in presence or absence of 5 mM of pyruvate. (C) Growth rate (h⁻¹) and lag time of PAO1 strain cultured in SCFM2 medium supplemented with 5 mM H₂O₂, in the presence or absence of 5 mM pyruvate. (D) Survival percentage following H₂O₂ exposure in SCFM2 medium for PAO1, *aceE*, and *aceF* mutant strains, and PAO1 supplemented with *aceF* strain’s supernatant (PAO1 (s)). (E) Pyruvate concentration (mM) in the supernatant of PAO1, *aceE*, and *aceF* mutant cultures after 4, 8, and 24 hours of incubation in SCFM2. In all panels, data are presented as mean ± SD. Each icon represents a biological replicate. Except for panel A, statistical significance was determined using one-way ANOVA with Tukey’s multiple comparisons test. Significance is indicated as * (p < 0.05), ** (p < 0.01), *** (p < 0.001), and **** (p < 0.0001). Source data are provided as a Source Data file.

To further evaluate this neutralizing function on primary macrophages, we performed cytokine profiling by multiplex flow cytometry after induction with H_2_O_2_ in presence or absence of pyruvate. While H_2_O_2_ exposure led to an increased secretion of IL-6, IL-8 and IL-10 cytokines, levels returned to baseline with pyruvate, confirming absence of stress under neutralizing conditions (Fig. 1B; one-way ANOVA with Tukey’s multiple comparisons test, p<0.05).

To further characterize pyruvate activity against H_2_O_2_ on the bacterial physiology, we measured the growth rate and lag phase of PAO1, *aceE* and *aceF* strains in SCFM2 in the presence or absence of H_2_O_2_ and pyruvate (Fig. 1C and Fig. S1A and S1B). H_2_O_2_ significantly impaired growth and extended the lag phase in all strains, while pyruvate restored both parameters (Fig. 1C and Fig. S1A and S1B; two-way ANOVA with Tukey’s multiple comparisons test, p<0.05). Similarly, in an H₂O₂ killing assay, *aceE* and *aceF* mutants showed higher survival than PAO1, respectively, while PAO1 supplemented with *aceF* supernatant showed comparable protection (Fig. 1D; one-way ANOVA with Tukey’s multiple comparisons test, p<0.05). This result is in agreement with the pyruvate secretion profile of the strains, where the *aceE* mutant showed transient secretion, while the *aceF* continuously accumulated pyruvate (Fig. 1E). Since pyruvate reacts with H₂O₂ to form acetate, which can complement PDHc mutations ^24,28^, we observed that acetate produced by the neutralizing reaction between pyruvate and H_2_O_2_ increased the growth rate of the *aceF* mutant but not that of PAO1 or *aceE* strain (Fig. S1C). A similar effect was seen in LB medium, which contains traces of acetate (Fig. S1D).

These results indicate that pyruvate protects against H_2_O_2_ oxidative stress both in *in vivo*-like infection environments and in laboratory condition.

### Reduced virulence and immune cells recruitment during airway infection

Metabolic crosstalk in host-pathogen interactions is a particularly fundamental process since the availability and competition for nutrients can influence the physiology of both infecting bacteria and the host. To assess whether pyruvate secretion by the *aceE* and *aceF* mutant strains changes the virulence potential of the infecting bacteria and the host response, we conducted infection assays using ALI infection models (Fig. 2).

**Figure 2.**
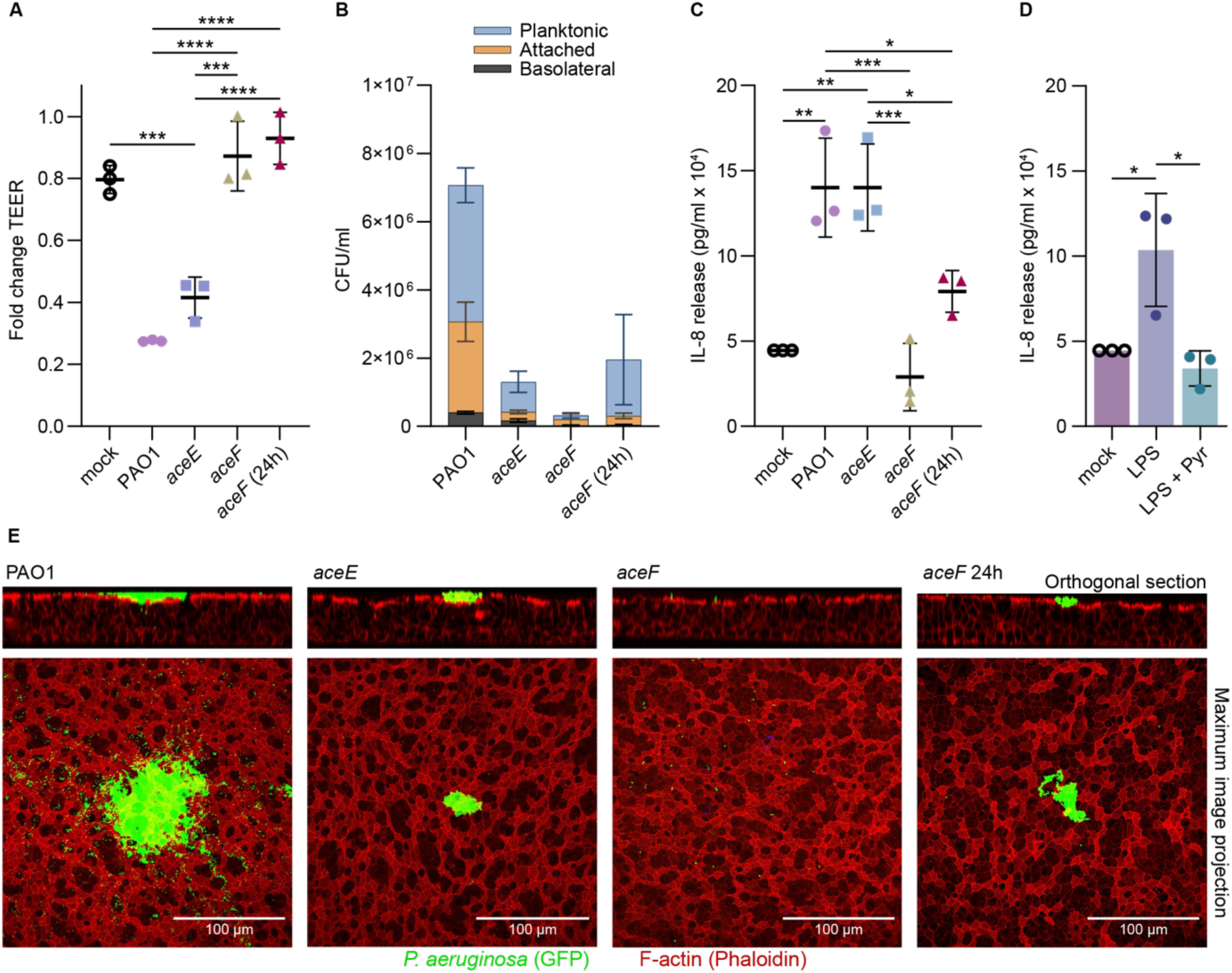
Mutations in *aceE* and *aceF* influence airway infection dynamics of *P. aeruginosa*. (A) Fold change in transepithelial electrical resistance (TEER), (B) bacterial counts (CFUs) for bacteria growing as planktonic cells in the apical compartment, those firmly attached to the epithelium, and those present in the basolateral compartment, and (C) Interleukin-8 (IL-8) secretion following 16 hours of infection with PAO1, *aceE*, and *aceF* mutant strains, and after 24 hours for the *aceF* strain, using an air-liquid interface (ALI) model of fully differentiated BCi-NS1.1 airway cells. (D) IL-8 secretion into the basolateral medium following stimulation with 10 μL of 100 μg/mL LPS and inhibition with 10 μL of 10 mM pyruvate. “Mock” refers to uninfected control cells. Statistical significance was determined using one-way ANOVA with Tukey’s multiple comparisons test for TEER and IL-8 quantification. Significance is indicated as * (p < 0.05), ** (p < 0.01), *** (p < 0.001), and **** (p < 0.0001). Data represent the mean ± SD and icons represent each biological replicate. Source data are provided as a Source Data file. (E) Confocal microscopy images of fully differentiated ALI models infected with *P. aeruginosa* PAO1, *aceE* or *aceF* for 16 hours and *aceF* for 24 hours, expressing GFP in green, and the epithelial actin stained with phalloidin in red. Scale bar = 100 *μ* m.

Since the *aceF* mutant strain has a reduced growth rate relative to PAO1 in laboratory conditions (Fig. S1B), we performed infection assays and quantified infection parameters after 16 hours of infection and additionally at 24 hours for the *aceF* mutant strain. The additional 24 hours time point for the *aceF* mutant was included to ensure that bacterial loads were comparable between strains and allow a more accurate comparison of infection parameters by minimizing confounding effects due to differences in growth kinetics. To quantify bacterial cytotoxicity and colonization capacity, we measured transepithelial electrical resistance (TEER), which reflects epithelial barrier integrity, and bacterial CFUs in the ALI infection models. CFUs were assessed for bacteria growing as planktonic cells in the apical compartment, those firmly attached to the epithelium, and those present in the basolateral compartment after penetration through the epithelium.

TEER measurements revealed a lower capacity of the *aceF* mutant strain to damage the epithelium after 24 hours compared to PAO1 after 16h of infection. In contrast, the *aceE* mutant strain induced a comparable epithelial damage to that of PAO1 (Fig. 2A), suggesting that this mutation does not impair bacterial cytotoxicity. However, the bacterial CFUs were significantly lower for both mutant strains relative to PAO1 indicating a clear fitness cost of these PDHc mutations in *in vivo*-like conditions (Fig. 2B). Additionally, the *aceF* mutant strain triggered significantly lower IL-8 secretion (Fig. 2C), a key cytokine involved in immune cell recruitment at the site of infection (two-way ANOVA with Tukey’s multiple comparisons test, p<0.05). Notably, extensive trans-epithelial penetration (CFUs in the basolateral side) was only observed for PAO1, whereas *aceE* and *aceF* mutant strains did not exhibit such an invasive behavior, potentially indicating a reduction in their virulence (Fig. 2B). These results were further supported by confocal microscopy analysis, which revealed distinct colonization patterns between PAO1, *aceE*, and *aceF* on the airway epithelial surfaces (Fig. 2E).

As the 24 hours infections with the *aceF* mutant strain exhibited similar bacterial growth to the 16 hours infection with the *aceE* mutant strain, but a significantly lower epithelial damage and IL-8 secretion, we hypothesized that the secreted pyruvate might also elicit immunosuppression activity on the epithelial cells. To investigate this hypothesis, we stimulated the epithelium by adding lipopolysaccharide (LPS), which elicits a strong immune response, and tested the effect of pyruvate on the ability of immune system recruitment by IL-8 quantification (Fig. 2D). Interestingly, we observed a significant reduction of IL-8 release in the presence of LPS + pyruvate as compared to LPS alone, supporting our hypothesis that pyruvate can additionally act as an immunosuppressor on the airway epithelium (Fig. 2D, one-way ANOVA with Tukey’s multiple comparisons test, p<0.05).

These results suggest that mutations in the *aceE* and *aceF* genes contribute to persistent infections and establish a potential link between pyruvate metabolism, virulence regulation and modulation of the immune system recruitment in *P. aeruginosa* infections.

### Pyruvate modulates macrophage killing capacity and limits cytokines release

Macrophages constitute one of the first lines of defense against airway infection. Since ALI infections revealed reduced IL-8 secretion upon infection with the *aceF* strain, we hypothesize that pyruvate secretion may modulate macrophage responses.

To test this hypothesis, we performed infection assays using human primary monocytes-derived macrophages and quantified both phagocytic bacterial killing and macrophage activation by cytokines release (Fig. 3). Killing assays revealed that *aceE* and *aceF* mutant strains were more resistant to intracellular killing with a large fraction of phagocytosed bacteria (51% to 74% survival) remaining viable within the macrophages after 4 hours of infection (Fig. 3A; one-way ANOVA with Tukey’s multiple comparisons test, p<0.05). In contrast, no PAO1 CFUs were detected at the same timepoint (0% survival), consistent with the already lower intracellular CFUs quantified following bacteria-macrophage incubation and gentamycin treatment at t0. Importantly, supplementation with pyruvate at the time of PAO1 infection led to an increased bacterial survival (23% survival), supporting pyruvate’s role in promoting survival to macrophage killing (Fig. 3A; one-way ANOVA with Tukey’s multiple comparisons test, p<0.05).

**Figure 3.**
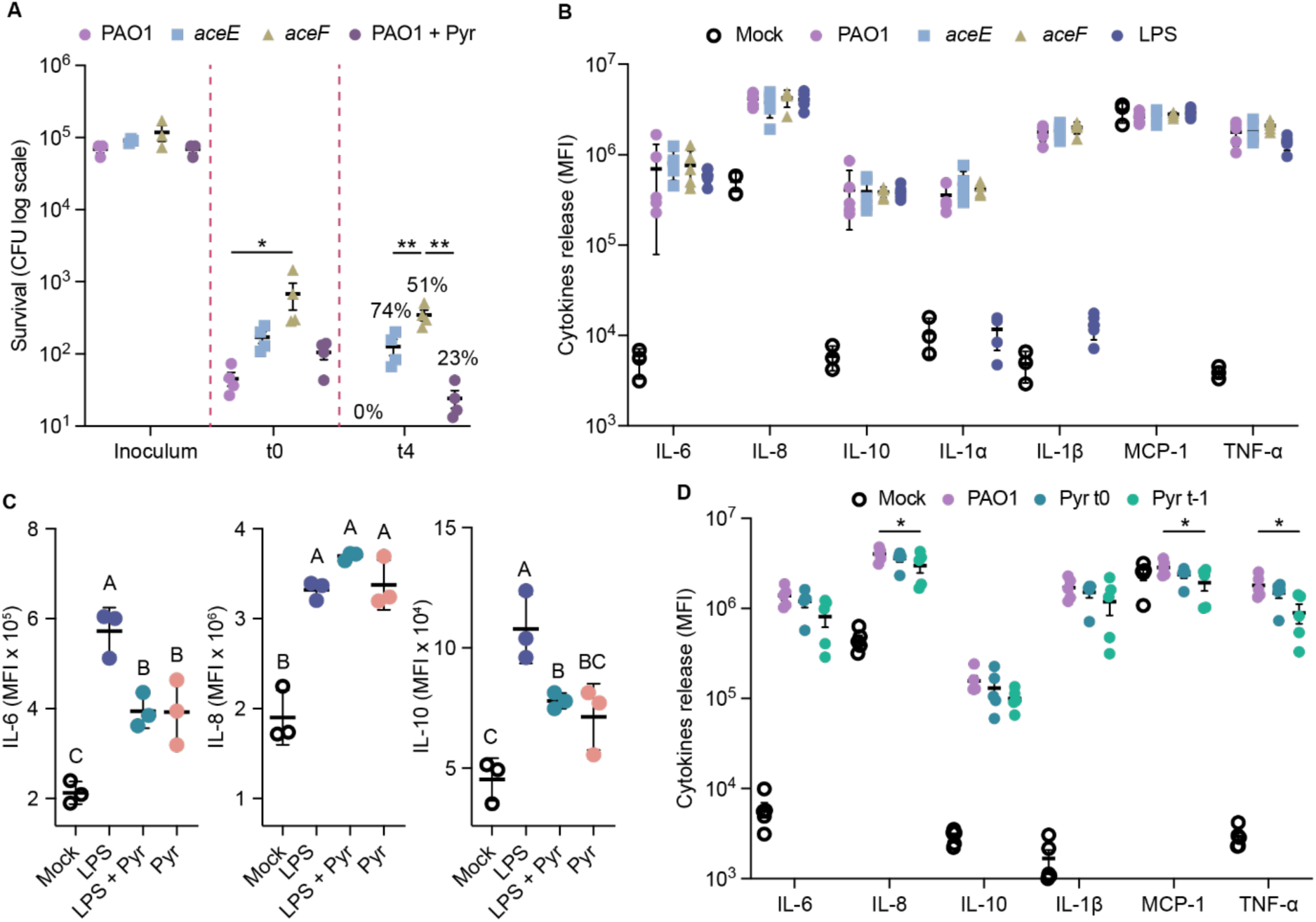
Pyruvate reduces macrophage activation and limits bacterial killing. (A) Intracellular bacterial survival upon macrophage phagocytosis of PAO1, *aceE* and *aceF* strains. For PAO1 + Pyr, 5 mM of pyruvate was added at the time of infection. Inoculum represent the bacterial count before infection, t0 represent the time point after 1 hour bacterial phagocytosis followed by 1 hour gentamycin treatment to remove extracellular bacteria. (B) Cytokines release upon infection with PAO1, *aceE* and *aceF* strains. LPS was used as positive control of macrophage stimulation. (C) IL-6, IL-8 and IL-10 release upon stimulation with LPS in presence and absence of 5 mM of pyruvate. (D) Cytokines release upon infection with PAO1 in presence or absence of 5 mM of pyruvate. Pyruvate was added either at the time of infection (t0) or macrophages were pre-incubated for 1 hour prior infection (t-1). For cytokine stimulation assays, data are presented as mean fluorescence intensity (MFI). Statistical significance was determined using one-way (A and C) or two-way (B and D) ANOVA with Tukey’s multiple comparisons test. Significance is indicated as * (p < 0.05), ** (p < 0.01) or by connecting letters (C). Data represent the mean ± SEM and icons represent each biological replicate. Source data are provided as a Source Data file.

Despite these differences in bacterial survival, macrophages infected with PAO1, *aceE*, or *aceF* for 4 hours showed comparable cytokine profiles for IL-6, IL-8, IL-10, IL-1α, IL-1β, MCP-1 and TNF-α (Fig. 3B; one-way ANOVA with Tukey’s multiple comparisons test, p>0.05). However, when assessing pyruvate secretion by the *aceE* and *aceF* strain under the infection conditions, we found that no pyruvate was produced within the short time frame of the experiment. To further evaluate the role of pyruvate in immune cell activation, we stimulated macrophages with LPS in the presence or absence of exogenous pyruvate. While LPS triggered a classic inflammatory response characterized by increased IL-6 and IL-10, pyruvate addition suppressed secretion of both cytokines (one-way ANOVA with Tukey’s multiple comparisons test, p<0.05), while no difference was instead observed for IL-8 (Fig. 3C; one-way ANOVA with Tukey’s multiple comparisons test, p>0.05). A similar result was obtained when infecting macrophages with PAO1 in presence of exogenous pyruvate either added at the time of infection (Pyr t0), or by pre-incubating macrophages for one hour with pyruvate (Pyr t-1), where pre-incubation decreased the release of IL-8, MCP-1 and TNF-α relative to untreated PAO1 infections (Fig. 3D; one-way ANOVA with Tukey’s multiple comparisons test, p<0.05). Other cytokines showed a similar decreasing trend, albeit the differences did not reach statistical significance.

Together, these results indicate that pyruvate secretion enhances bacterial survival within macrophages and dampens macrophage activation by reducing the release of key pro-inflammatory cytokines.

### Clinical relevance of PDHc mutations

To assess the consequences of PDHc mutations and their role in metabolic cross-talk during infection, we selected from our longitudinal collection of *P. aeruginosa* clinical isolates from pwCF having persistent infections, 12 clinical strains each containing a different non-synonymous mutation in the PDHc encoding genes *aceE* and *aceF* ^29^. The selected strains represent diverse clone types from different pwCF and are representative of the 89 isolates in our collection containing PDHc mutations (Fig. 4A) ^29^. To determine whether these strains exhibit a similar phenotype to the recombinant *aceE* and *aceF* mutant strains, we measured their maximum growth rate and pyruvate secretion during the growth in SCFM2 medium. The growth rate showed a significant variability, with a range from 0.321 to 1.234 indicating a broad fitness cost of the mutations (Fig. 4B). Additionally, all clinical strains secreted high amount of pyruvate during growth (Fig. 4C), with concentrations comparable to those of the recombinant strains and previously analyzed clinical strains ^24,30^. As observed for the *aceE* and *aceF* mutant strains, two distinct patterns of pyruvate secretion were observed: one involving secretion followed by re-assimilation, and another one characterized by continuous secretion (Fig. 4C). The relationship between the secretion patterns and the type or location of mutations remains unclear, as both point mutations and insertion-deletion events occur in the *aceE* and *aceF* genes, independently of secretion pattern. However, it confirms the extent of the metabolic crosstalk between *P. aeruginosa* and the host in different infection scenarios.

**Figure 4.**
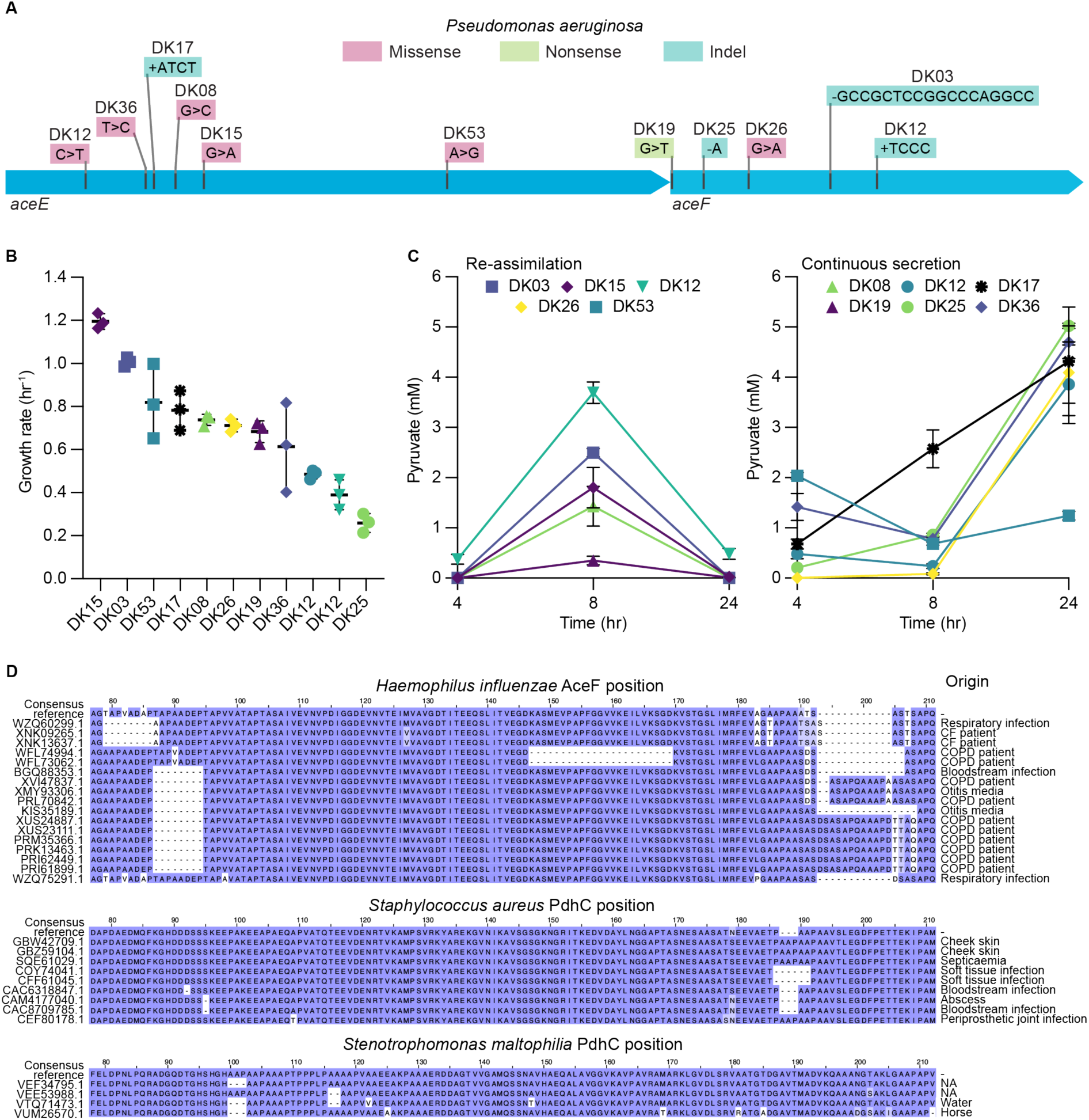
Characterization of clinical strains with mutations in the PDHc. (A) Type of non-synonymous mutations observed in the *aceEF* operon in different clinical isolates from chronically infected pwCF belonging to different clone types. (B) Growth rate (hour^−1^) in SCFM2 of clinical strains harboring PDHc mutations colored by their clone type. Data represent the mean ± SD and icons represent each biological replicate. (C) Pyruvate secretion (mM) of clinical strains harboring PDHc mutations over 24 hours of growth in SCFM2. The distinct patterns are indicated by re-assimilation and continuous secretion. Data represent the mean ± SEM of three independent biological replicates. Source data are provided as a Source Data file. (D) Multiple sequence alignments of PDHc subunit 2 proteins from *Haemophilus influenzae*, *Staphylococcus aureus*, and *Stenotrophomonas maltophilia*. Sequences were selected based on the presence of high-impact mutations and traceable clinical metadata, except for *S. maltophilia*, for which all sequences harboring high-impact mutations were included regardless of metadata availability.

Since pyruvate is a central metabolite conserved across diverse species, we hypothesized that similar mutations in the PDHc might also arise in other pathogens associated with persistent infections. To test this possibility, we aligned all complete sequences of the PDHc subunits 1 and 2 from *S. aureus*, *H. influenzae*, *S. maltophilia* and *P. aeruginosa* available on the National Center for Biotechnology Information (NCBI) database. We observed greater variability in subunit 2, particularly in the form of high-impact mutations (insertions, deletions, or frameshifts), which were frequently detected in strains with metadata confirming a clinical origin, both in *P. aeruginosa* (Supplementary Figure 2) and in the other species (Fig. 4D). In contrast, high-impact mutations in subunit 1 were less common, and only some *S. aureus* strains with such mutations could be traced back to clinical isolates (Supplementary Figure 2). It is important to note that additional functionally relevant PDHc mutations may exist in clinical strains, since our targeted analysis was restricted only to clearly defined high-impact variants associated with clinical metadata. Overall, these findings indicate that PDHc mutations are not restricted to *P. aeruginosa* but are also present in clinical isolates of other prevalent airway pathogens.

## Discussion

Metabolism is a fundamental process dynamically regulated to meet the nutrient availability and physiological demands of an organism. In *P. aeruginosa*, clinical isolates have been shown to undergo metabolic specialization during infection, suggesting adaptive responses to the host environment ^9,30,31^. These shifts not only compensate for the fitness costs of other phenotypic traits but can also be actively selected to enhance bacterial persistence ^24^.

In this study, we investigated mutations in the *aceE* and *aceF* genes, encoding components of the PDHc, as a non-conventional persistence mechanism during chronic *P. aeruginosa* airway infection ^24,29^. These mutations prevent efficient processing of pyruvate, revealing that secreted pyruvate functions not just as a metabolic byproduct but as a bacterial immunometabolite that reshapes the host environment to favor persistence. We demonstrate that pyruvate exerts several functions at the host-pathogen interface (Fig. 5). First, it scavenges ROS through a non-enzymatic reaction, potentially reducing oxidative stress in the airway microenvironment. Second, it reduces the host immune response by limiting IL-8 production and, therefore, recruitment of immune cells. Third, it limits macrophage activation and enhances bacterial survival after phagocytosis. Chronic activation of innate immune cells and epithelial dysfunction in infected CF airways generate sustained ROS levels ^32^. While the antioxidant roles of pyruvate have been well described in mammalian systems ^14–16^, in this case, pyruvate secretion by the pathogen attenuates oxidative killing and inflammation, effectively hijacking a metabolic circuit used by the host to protect its own tissues. Notably, the reaction of pyruvate with H₂O₂ yields acetate, which may be re-assimilated by PDHc mutant strains and shuttled into the TCA cycle, partially compensating for their metabolic deficit ^28^.

**Figure 5.**
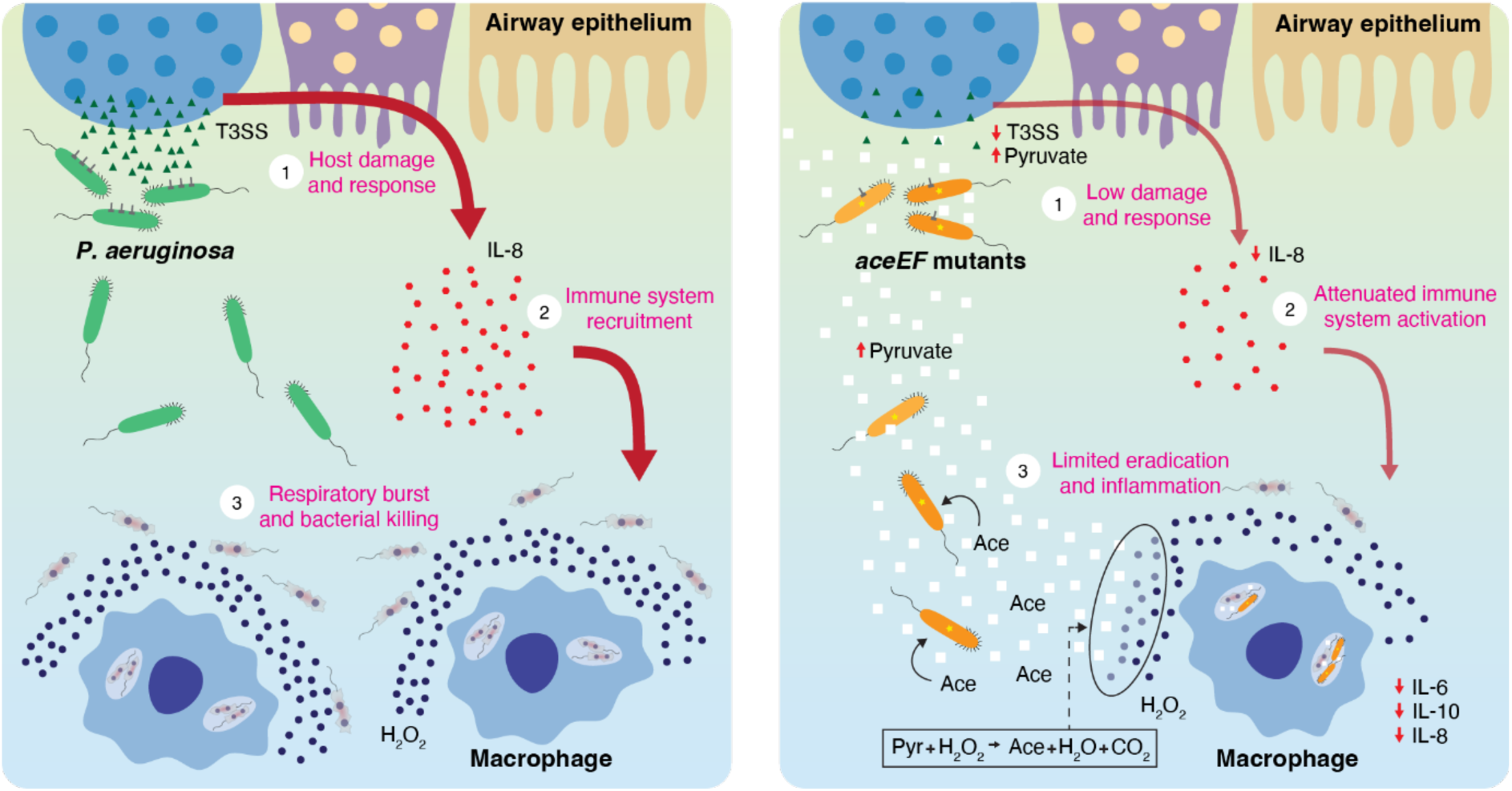
Persistence mechanism through PDHc mutations in *P. aeruginosa*. (A) Infection dynamics of wild type strains: 1) the secretion of T3SS causes damage to the airway epithelium and consequently elicits a host response; 2) the host secretes IL-8 for immune system recruitment; 3) macrophages reach the infection site and produce ROS, including H_2_O_2_, to eradicate the extracellular and intracellular (phagolysosome) infecting bacteria. (B) Infection dynamics of the *aceE* and *aceF* mutant strains: 1) reduced T3SS activity and the secretion of pyruvate lead to a reduced damage and host response; 2) low levels of secreted IL-8 by epithelial cells and of secreted IL-6, IL-8 and IL-10 by macrophages cause an attenuated immune response; 3) fewer innate immune cells arrive at the infection site to produce ROS which is neutralized by the pyruvate present in the environment; intracellular bacteria increase their survival within the phagolysosomes by pyruvate release; acetate is produced as by-product of the pyruvate-H_2_O_2_ reaction which can be assimilated by PDHc mutant strains compensating for their low growth rate.

Oxidative stress within the host environment may, therefore, represent a key selective pressure driving the emergence of *aceEF* mutations. However, when *P. aeruginosa* has experimentally evolved under oxidative stress in laboratory conditions, the predominant adaptive response involves mutations that lead to overexpression of catalases ^33^. While pyruvate reacts non-enzymatically with H_2_O_2_, catalases achieve a similar antioxidant outcome through an enzymatic reaction ^34^. This difference in selection dynamics suggests that, although both mechanisms reduce oxidative burden, *aceEF* mutations may not merely reflect a response to oxidative pressure, but rather a broader adaptation to the host environment. Consistent with this result, our findings show that secreted pyruvate not only scavenges ROS, but also reduces IL-8 secretion by airway epithelial cells, dampens macrophage activation, and promotes bacterial survival after phagocytosis. Importantly, this effect is independent of the downregulation of classical virulence pathways. As previously described, PDHc mutants exhibit hallmarks of chronic infection, including reduced expression of the T3SS, upregulation of T6SS or increased production of phenazines and alginate ^7,24,35^. While T3SS suppression likely contributes to reduced host immune activation since its effectors can induce inflammatory responses ^36^, our data show that pyruvate alone can reduce cytokine production.

Immune activation requires a profound metabolic reprogramming that both supports and defines effector functions ^37^. Pyruvate lies at the core of this process, as its metabolic fate determines the balance between glycolytic and oxidative metabolism. In this context, exogenous pyruvate, here produced by infecting bacteria, can perturb this delicate equilibrium, disrupting the metabolic reprogramming required for immune activation and thereby dampening the host immune response ^38,39^. This immunomodulatory effect of pyruvate may contribute to a form of disease tolerance, in which host tissue damage and inflammation are reduced despite the persistence of the pathogen. Our findings conceptually parallel the previously described itaconate–alginate interplay in *P. aeruginosa*, which helps limit immunopathology ^4,11^. However, our study extends this paradigm by identifying pyruvate as a universally conserved and central metabolite with immune modulator function. Unlike alginate, which is species-specific, pyruvate can be secreted by a wide range of bacteria under metabolic stress or imbalance. This observation raises the possibility that pyruvate immune modulation is a strategy for promoting chronic colonization and tissue tolerance in other pathogens as well ^20,21^. Indeed, we identified mutations in the PDHc in clinical strains from infection sites in *S. aureus*, *H. influenzae* and *S. maltophilia*, supporting the notion that modifications in pyruvate metabolism constitute a recurrent feature of pathogen adaptation during infection. Moreover, *in-situ* analyses of sputum samples from pwCF showed a decrease in expression of the PDHc indicating that similar effect could potentially also be achieved through regulatory mechanisms ^40^. Given its dual role in metabolism and signaling, pyruvate may have broader and underappreciated effects on immune regulation beyond airway infections ^41^.

More broadly, our work emphasizes the emerging role of secreted metabolites as signaling molecules in host-pathogen interactions. Future studies should explore whether pyruvate is part of a broader repertoire of metabolites with immunomodulatory effect. It will also be important to assess whether interfering with pyruvate-mediated signaling, by limiting its secretion, counteracting its effects or restoring host redox balance, could offer novel strategies for managing chronic infections without relying solely on antimicrobial therapies. Ultimately, targeting such metabolic pathways may not only impair bacterial persistence but also modulate the host’s tolerance to infection, providing a complementary approach to conventional antimicrobial strategies.

## Materials and Methods

### Strains and media

The *aceE* and *aceF* mutant strains in a *P. aeruginosa* PAO1 background used in this study were previously generated in ^24^ and clinical strains used in this work were previously described in ^29^. Bacteria were grown in SCFM2 medium prepared as previously described in ^42^, excluding DNA and mucins to reduce viscosity, or in LB medium. Cultures were grown at 37°C and 250 rpm.

### Bacterial growth assays

Bacterial growth was assessed at 37°C and 250 rpm in a BioTek ELx808 Absorbance Microtiter Reader (BioTek Instruments, Winooski, Vermont, USA) by measuring OD_600nm_ every 20 minutes for 24 hours. Growth was initiated by inoculating an overnight bacterial culture in LB or SCFM2 media supplemented with either 5 mM H_2_O_2_, 5 mM pyruvate, 5 mM acetate or the pre-incubated reaction at 37°C and 250 rpm for 30 minutes of 5 mM H_2_O_2_ with 5 mM pyruvate.

### Pyruvate concentration determination

Bacterial cultures were initiated at OD_600nm_=0.01 in SCFM2. The supernatants were collected by centrifugation at 5.000 x *g* for 10 minutes at 4, 8 and 24 hours after inoculation. For the quantification of pyruvate in the respective supernatants, the Pyruvate Assay Kit (Sigma-Aldrich) was used following the manufacturer’s instructions.

### H_2_O_2_ killing assays

H_2_O_2_ killing assays were performed as previously described ^43^, with minor modifications. Late exponential phase cultures (OD_600nm_≈2.5) in SCFM2 were normalized with the supernatant of an overnight culture of the same strain to an OD_600nm_ of 0.5. 5 mM H_2_O_2_, or 5 mM H_2_O_2_ together with 5mM of pyruvate, were incubated with the normalized cultures. After 30 minutes, 0.2% (w/v) sodium thiosulfate was added to neutralize residual H_2_O_2_. CFUs were determined by the standard microdilution technique on LB agar plates. Survival was reported as percentage of CFUs counted in the treated sample relative to the CFUs counted in the samples before treatment.

### Air-Liquid Interface infection model preparation

BCi-NS1.1 cells, a human airway basal cell line with immortalized properties and the ability to differentiate into secretory, goblet, Clara, and ciliated cells under ALI conditions ^44^, were utilized to establish ALI infection models, as previously described ^45,46^. Initially, the cells were expanded in culture flasks maintained at 37 °C with 5% CO_2_ in a humidified incubator, using Pneumacult-Ex Plus medium (STEMCELL Technologies) as per the manufacturer’s guidelines. Following expansion, 10^5^ cells were seeded onto polyester membrane inserts (1 μm pore size, Falcon) pre-coated with type I collagen (Gibco). Once confluency was achieved, ALI conditions were induced by removing the medium from the apical chamber and replacing the basolateral medium with Pneumacult-ALI maintenance medium (STEMCELL Technologies), supplemented with 4 μg/mL heparin, 480 ng/mL hydrocortisone, Pneumacult-ALI supplement, and Pneumacult-ALI maintenance supplement (STEMCELL Technologies). The ALI cultures were maintained at 37 °C and 5% CO_2_ in a humidified incubator for 28 days, with media replenished every 3-4 days. TEER was routinely monitored using a chopstick electrode (STX2; World Precision Instruments) to assess epithelial polarization. After 15 days of ALI culture, accumulated mucus was cleared by washing the apical surface with PBS every seven days.

### Epithelial infection procedure and characterization

A 10 μL drop of diluted overnight culture, containing 10^3^ CFUs, was applied to the apical surface of fully differentiated BCi-NS1.1 ALI cultures, with 600 μL of supplemented Pneumacult-ALI maintenance medium present in the basolateral chamber. Control wells received an equivalent volume of bacteria-free PBS. The infected cells were incubated at 37 °C with 5% CO_2_ in a humidified incubator for the required duration to assess the infection. After incubation, 200 μL of PBS was added to the apical surface, and TEER measurements were taken as described above. The 200 μL of PBS from the apical side and the 600 μL of medium from the basolateral chamber were collected, and bacterial CFUs in these samples were quantified by plating 10 μL of serial dilutions onto LB agar plates. The remaining samples were stored at -80 °C for future analysis. At this stage, the cells were either prepared for bacterial CFU counting on the epithelium or for imaging via confocal microscopy. For CFU quantification, 200 μL of PBS was added to the apical surface, and the epithelial layer was disrupted using a cell scraper. The resulting suspension was then subjected to LB agar plating of 10 μL in serial dilutions to determine CFU counts.

To prepare infected ALI cultures for confocal microscopy, samples were rinsed once with PBS and fixed with 4% paraformaldehyde, which was added to both the apical and basolateral chambers, for 20 minutes at 4 °C. Following fixation, cells were washed three times with PBS, then permeabilized and blocked by incubating them in a buffer containing 3% BSA, 1% Saponin, and 1% Triton X-100 in PBS for 1 hour at room temperature. A 100 μL staining solution containing Phalloidin-AF555 (Invitrogen) diluted at 1:400 in a staining buffer (3% BSA and 1% Saponin in PBS), was applied to the apical chamber. The cells were then incubated for 2 hours at room temperature for staining. Fixed and stained epithelial layers were carefully detached from their supports using a scalpel and mounted onto glass slides with VECTASHIELD Antifade Mounting Medium (VWR). Imaging of infected cultures was performed using a Leica Stellaris 8 Confocal Microscope (40x magnification, 1.3 oil) and analyzed with LasX software 1.4.4.26810 (Leica).

The basolateral media collected at the end of the infection was also used to measure IL-8 release, following the manufacturer’s protocol for the Human IL-8/CXCL8 DuoSet ELISA Kit (R&D Systems).

### H_2_O_2_ killing assays in ALI cultures

To assess the effectiveness of pyruvate in protecting bacteria colonizing the epithelial infection system from H_2_O_2_, the infection in fully differentiated ALI models was first established as previously described. After 6 hours of incubation at 37 °C and 5% CO_2_ in a humidified incubator, 10 μl of 2 mM H_2_O_2_ or first 10 μl of 5 mM pyruvate was added to the apical chamber followed by 10 μl of 2 mM H_2_O_2_. The system was then further incubated at 37 °C and 5% CO_2_ for an additional 30 minutes. Subsequently, 0.2% (w/v) sodium thiosulfate was added to neutralize residual H_2_O_2_, TEER was measured and CFUs present in the apical chamber, basolateral medium, or attached to the epithelium were quantified by plating 10 μL of serial dilutions of respective bacterial suspensions on LB agar plates, as previously described. Finally, the percentage of bacterial survival was calculated by comparing CFU counts with those of untreated epithelial cultures.

### Determination of epithelial IL-8 release inhibition by pyruvate

In order to induce the release of IL-8 by the epithelium, 1000 ng of LPS with or without 10 μl of 10 mM pyruvate were inoculated into the apical surface of fully differentiated ALI models. The basolateral media was collected after 16 hours and IL-8 release was measured in those samples following the manufacturer’s protocol for the Human IL-8/CXCL8 Duoset ELISA Kit (R&D Systems).

### Macrophages preparation

Peripheral blood mononuclear cells (PBMCs) were isolated from fresh buffy coats obtained from healthy donors through the Department of Clinical Immunology, Blood Bank, at Rigshospitalet (Copenhagen, Denmark). This study was approved by the local ethics committee of the Capital Region of Denmark (Region Hovedstaden), registration number H-20024750. Buffy coats from 3–4 donors were pooled for each experiment to increase cell yield and reduce donor-specific variability. PBMCs were processed on the day of collection. 50 mL of buffy coat were diluted 1:1 in PBS supplemented with 2% fetal bovine serum (FBS; Gibco). PBMCs were isolated using 50 mL LEUCOSEP tubes (Greiner) pre-filled with Lymphoprep (STEMCELL Technologies). After isolation, cells were washed twice in PBS + 2% FBS + 2 mM EDTA and centrifuged at 120 × g for 10 minutes at 4 °C to remove residual platelets. PBMCs were then cryopreserved at 1 × 10⁷ cells/mL in FBS containing 10% dimethyl sulfoxide (DMSO; Thermo Fisher Scientific).

Cryopreserved PBMCs were rapidly thawed at 37 °C and immediately transferred into pre-warmed M0 base medium consisting of RPMI 1640 (Gibco) supplemented with 10% FBS and 50 ng/mL macrophage colony-stimulating factor (M-CSF; Miltenyi Biotec). Monocytes were purified by negative selection using the MojoSort™ Human Pan Monocyte Isolation Kit (BioLegend), according to the manufacturer’s instructions. Purified monocytes were seeded at 1.4 × 10⁷ cells/mL in M0 medium in P-100 tissue culture-treated dishes and maintained at 37 °C and 5% CO₂. Cells were differentiated into macrophages over 6 days, with a medium change at 24 hours to remove non-adherent cells that were seeded in a new dish.

For macrophage harvesting, cultures were incubated with TrypLE (Gibco) for 5–10 minutes at 37 °C, cells were detached using a cell scraper, and TrypLE was neutralized with cold PBS. Macrophages were pelleted at 300 × g for 5 min and resuspended at the desired concentration. A total of 1.25 × 10⁵ cells were seeded per well in 48-well clear flat-bottom tissue culture-treated plates (Falcon) and incubated for 24 hours before downstream assays.

### Macrophages induction assay

Before each experiment, the supernatant from macrophages seeded in 48-well plates was removed and replaced with 300 μL of fresh M0 medium. Macrophages were then stimulated with *P. aeruginosa* at a multiplicity of infection (MOI) of 1, H₂O₂ (5 mM), LPS (1 μg/mL), or the same conditions supplemented with pyruvate (5 mM), as indicated. After 4 hours at 37 °C and 5% CO₂, supernatants were collected and stored at −80 °C. Cytokine levels were quantified using the Cytometric Bead Array Flex Set (CBA; BD Biosciences) for human IL-6, IL-8, IL-10, IL-1β, IL-1α, MCP-1 and TNF-α, in a NovoCyte Quanteon Flow Cytometer (Agilent) following the manufacturer’s instructions. Three to five biological replicates were analyzed, each in technical triplicate, and a non-treated group served as the mock control.

### Phagocytosis and intracellular survival assay

Before infection, the supernatant in each well was removed and replaced with 300 μL of fresh M0 medium. Macrophages were infected at a MOI of 1 with selected *P. aeruginosa* strains prepared from overnight cultures. The inoculum dose was verified by serial plating on LB agar. Immediately after infection, plates were centrifuged at 1,200 × g for 5 minutes to promote bacterial contact with macrophages. To eliminate extracellular bacteria, gentamicin was added at 1 hour post-infection (hpi) to a final concentration of 50 μg/mL. For quantification of phagocytosis, cells were washed with PBS at 2 hpi and lysed in PBS containing 0.5% (v/v) Triton X-100. CFUs were determined by serial dilution and plating, representing the number of internalized bacteria.

Intracellular survival was assessed using a parallel plate infected with the same inoculum and macrophage batch. At 6 hpi, cells were washed with PBS and lysed in PBS + 0.5% (v/v) Triton X-100. CFUs obtained from serial dilutions reflected bacterial survival within macrophages after 6 h.

### Protein alignments

Protein sequences corresponding to PDHc subunits 1 and 2 from *P. aeruginosa*, *S. aureus*, *H. influenzae*, and *S. maltophilia* were retrieved from the National Center for Biotechnology Information (NCBI) protein database, by the query “((Pseudomonas aeruginosa[Organism]) AND aceE) NOT “partial”[Title]”, “((Pseudomonas aeruginosa[Organism]) AND aceF) NOT “partial”[Title]”, “((Staphylococcus aureus[Organism]) AND pdhA) NOT “partial”[Title]”, “((Staphylococcus aureus[Organism]) AND pdhC) NOT “partial”[Title]”, “((Haemophilus influenzae[Organism]) AND aceE) NOT “partial”[Title]”, “((Haemophilus influenzae[Organism]) AND aceF) NOT “partial”[Title]”, “((Stenotrophomonas maltophilia[Organism]) AND pdhA) NOT “partial”[Title]”, “((Stenotrophomonas maltophilia[Organism]) AND pdhC) NOT “partial”[Title]”, respectively. Entries annotated as partial were excluded from the analysis. Multiple sequence alignments were performed using MAFFT software (v7) ^47^ with default parameters.

### Quantification and statistical analyses

Growth rates and maxOD were calculated in the QurvE 1.1 software. Data were analyzed using GraphPad Prism 10.3.1 software (GraphPad Software). Statistically significant differences were examined using ordinary one-way ANOVA or two-way ANOVA test with Tukey’s multiple comparisons test to derive the significance of differences between all groups of each experiment. A p value of <0.05 was considered statistically significant.

### Data availability

The authors declare that all data necessary for suppor2ng the findings of this study are enclosed in this paper (Source Data).

## Acknowledgements

We thank the Infection Microbiology group for their insightful comments and discussion, especially Alexander Frederick Melanson for his support with macrophage assays. The Basal Cell Immortalized Non-Smoker 1.1 (BCi-NS1.1) cell line was a gift from Professor Ronald G. Cristal (Weil Cornell Medical College, New York, USA). This research was funded by the “Novo Nordisk Foundation Center for Biosustainability (CfB)” grant number NNF10CC1016517, by the Challenge Grant NNF19OC0056411 from The Novo Nordisk Foundation, and a grant from The John and Birthe Meyer Foundation to H.K.J. P.L. is the recipient of an ERS/EU RESPIRE4 Marie Skłodowska-Curie Postdoctoral Research Fellowship (Ref. nr: R4202305-01047; this project has received funding from the European Respiratory Society and the European Union’s H2020 research and innovation programme under the Marie Skłodowska-Curie grant agreement No 847462).

## Author’s contribution

RLR and PL designed the project. SL, LM and PL performed the experiments. RLR performed the data analysis. HKJ provided strains and clinical data. HKJ and SM provided funding. PL and RLR verified the data and wrote the first draft of the paper. All authors approved the final version of the manuscript.

